# Shared expression of Crassulacean acid metabolism (CAM) genes predates the origin of CAM in the genus *Yucca*

**DOI:** 10.1101/371344

**Authors:** Karolina Heyduk, Jeremy N. Ray, Saaravanaraj Ayyampalayam, Nida Moledina, Anne Borland, Scott A. Harding, Chung-Jui Tsai, Jim Leebens Mack

**Affiliations:** Department of Plant Biology, University of Georgia, Athens, Georgia, USA; Georgia Advanced Computing Resource Center, University of Georgia, Athens, Georgia, USA; School of Natural and Environmental Sciences, Newcastle University, Newcastle, United Kingdom; Department of Genetics, University of Georgia, Athens, Georgia, USA; Warnell School of Forestry, University of Georgia, Athens, Georgia, USA

**Keywords:** antioxidant, carbohydrates, hybrid, metabolomics, photosynthesis, RNA-seq, transcriptome, Yucca

## Abstract

**Highlight:** Although large differences in metabolism exist between C_3_ and CAM species, we find that many CAM genes have shared expression patterns regardless of photosynthetic pathway, suggesting ancestral propensity for CAM.

**Abstract:** Crassulacean acid metabolism (CAM) is a carbon-concentrating mechanism that has evolved numerous times across flowering plants and is thought to be an adaptation to water limited environments. CAM has been investigated from physiological and biochemical perspectives, but little is known about how plants evolve from C_3_ to CAM at the genetic or metabolic level. Here we take a comparative approach in analyzing time-course data of C_3_, CAM, and C_3_+CAM intermediate *Yucca* (Asparagaceae) species. RNA samples were collected over a 24-hour period from both well-watered and drought-stressed plants and were clustered based on time-dependent expression patterns. Metabolomic data reveals differences in carbohydrate metabolism and antioxidant response between the CAM and C_3_ species, suggesting changes to metabolic pathways are important for CAM evolution and function. However, all three species share expression profiles of canonical CAM pathway genes, regardless of photosynthetic pathway. Despite differences in transcript and metabolite profiles between the C_3_ and CAM species, shared time-structured expression of CAM genes in both CAM and C_3_ *Yucca* species suggests ancestral expression patterns required for CAM may have predated its origin in *Yucca*.

## Introduction

Crassulacean acid metabolism (CAM) is a carbon concentrating mechanism that can reduce photorespiration and enhance water use efficiency relative to plants that rely solely on the C_3_ photosynthetic pathway. In CAM plants, stomata are open for gas exchange at night, when transpiration rates are lower, and incoming CO_2_ is initially fixed by phosphoenolpyruvate carboxylase (PEPC) rather than RuBisCO. Carbon is temporarily stored as malic acid within the vacuole, and during the day stomata close and the malic acid is decarboxylated in the cytosol, ultimately resulting in high concentrations of CO_2_ around RuBisCO. The extra steps of CAM – carboxylation of PEP, decarboxylation of malic acid, transport into and out of the vacuole – come with extra energetic costs relative to C_3_ photosynthesis, but CAM plants have the advantage of acquiring carbon with increased water use efficiency (WUE). In addition, RuBisCO is able to act more efficiently with a high concentration of CO_2_ and the risk of photorespiration is thought to be significantly minimized (Cushman and Bohnert, 1997; Schulze *et al.*, 2013). CAM plants are therefore adapted to habitats where water stress is unavoidable and where the energetic cost of CAM is offset by reduced photorespiration under water limited conditions. CAM has evolved at least 35 independent times in flowering plants (Silvera *et al.*, 2010), thus making it a remarkable example of convergent evolution of a complex trait.

CAM has been studied from a biochemical standpoint for decades, and much is known about the carbohydrate turnover, starch cycling, and enzymatic machinery of CAM plants (Cushman and Bohnert, 1997; Chen *et al.*, 2002; Dodd *et al.*, 2002). Additionally, physiological studies of CAM plants have revealed the importance of succulence and large cells (Kluge and Ting, 1978; Nelson *et al.*, 2005; Nelson and Sage, 2008; Zambrano *et al.*, 2014). In terms of the basic machinery required for CAM, carbonic anhydrase (CA) aids in the conversion of CO_2_ to HCO_3-_ at night (Fig. 1A). PEPC fixes the carbon from CA into oxaloacetate (OAA), but its activity is regulated by a dedicated kinase, PEPC kinase (PPCK). Phosphorylated PEPC is able to fix carbon in the presence of its downstream product, malate, whereas the un-phosphorylated form is sensitive to malate (Nimmo, 2000; Taybi *et al.*, 2000). As day approaches, PPCK is down-regulated by a combination of two mechanisms: circadian regulation (Carter *et al.*, 1991; Hartwell *et al.*, 1996) or through metabolite control of transcription which results from elevation of cytosolic malate(Borland *et al.*, 1999). During the day, the stored malic acid exits the vacuole and is decarboxylated by either phosphoenolpyruvate carboxykinase (PEPCK) and/or NADP/NAD-malic enzyme, depending on CAM species (Holtum and Osmond, 1981). NADP/NAD-ME CAM plants additionally have high levels of PPDK, which converts pyruvate to PEP. This final step is important for CAM plants, as can be further used to generate carbohydrates and is also required for CO_2_ fixation during the following night.

**Figure 1.**
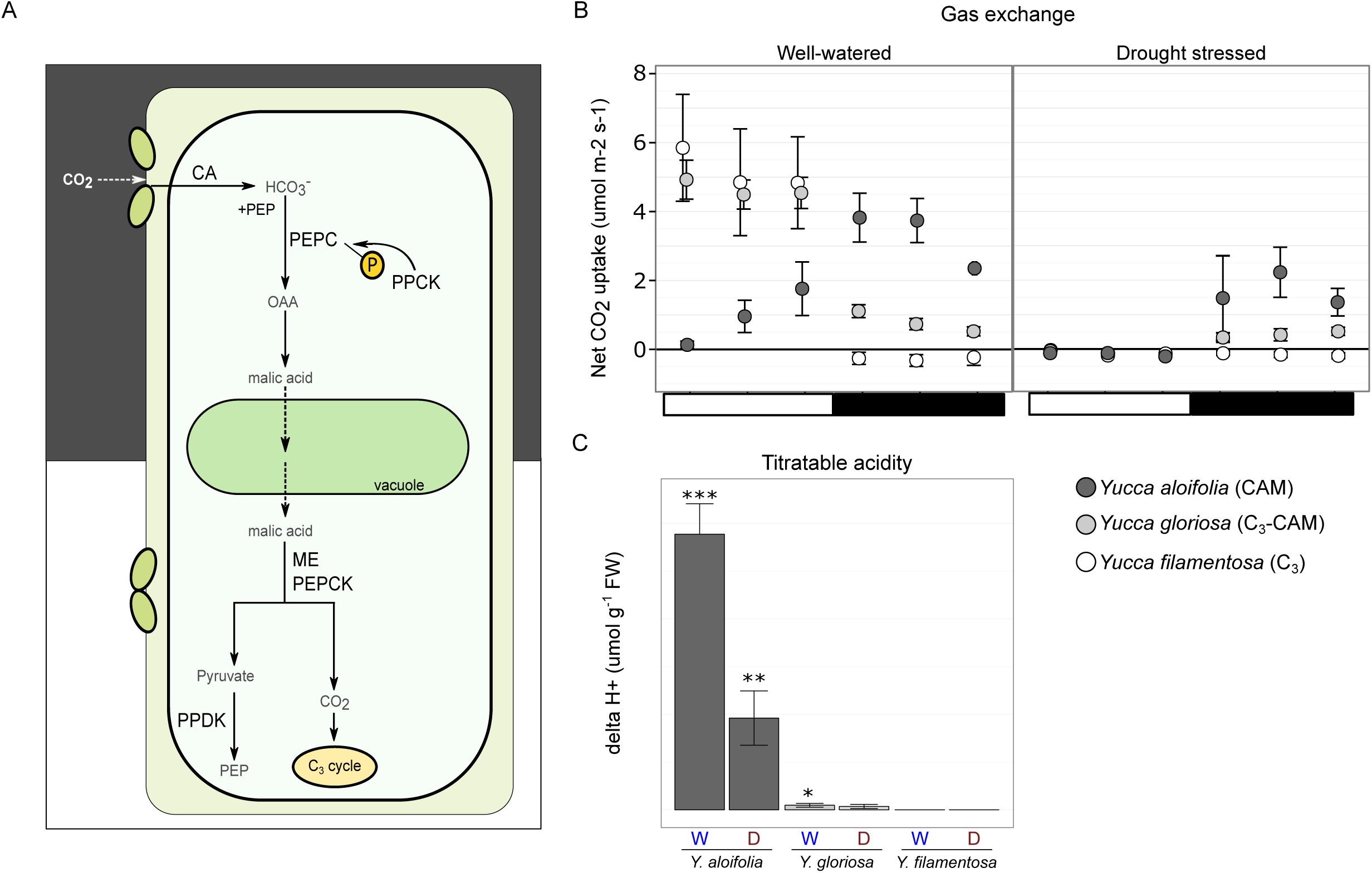
Overview of the CAM photosynthetic pathway and physiology of Yucca hybrid system. A) CAM pathway diagram. CA, carbonic anhydrase; PEP, phosphoenolpyuvate; PEPC, PEP carboxylase; PPCK, PEPC kinase; OAA, oxaloacetate; ME, malic enzyme; PEPCK, PEPC carboxykinase; PPDK, orthophosphate dikinase. B) Net CO2 accumulation on the same samples used for RNA-seq, with error bars representing 1 standard deviation from the mean. White bar and black bar under plots represent day and night, respectively. C) Delta H+ (the total titratable acid accumulated during the night) measured on samples used for RNA-seq from well-watered (“W”) and drought-stressed conditions (“D”), with error bars representing one standard deviation from the mean. Both gas exchange and titratable acidity plots are modified from data published in (Heyduk et al., 2016).

A daily turnover of sugars or starch for PEP generation is a defining characteristic of CAM plants. Carbohydrates that are laid down during the day must be broken down to PEP at night to provide substrate for CO_2_ fixation via PEPC. The nocturnal demand for PEP represents a significant sink for carbohydrates which CAM plants must balance with partitioning of carbohydrates for growth and maintenance (Borland *et al.*, 2016). The interplay between carbohydrate metabolism and CAM is clearly an important regulatory mechanism; previous work has shown that plants with reduced carbohydrate degradation have decreased CAM function at night (Dodd *et al.*, 2003; Cushman *et al.*, 2008*a*). The evolution of temporally integrated carbon metabolism in CAM plants presumably involves rewiring of gene regulatory networks to link these processes with the circadian clock. Although timing of photosynthetic gene expression is to some degree circadian controlled in both C_3_ and CAM species, the links between metabolism genes in CAM and circadian clock oscillators may be stronger (Hartwell, 2005; Dever *et al.*, 2015). CAM is typically described as a complex phenotype largely because of its role in central metabolism, and therefore its recurrent evolution is considered a remarkable and textbook example of convergence. Yet the frequent transitions to CAM across plants suggests that the evolutionary trajectory from C_3_ to CAM may not be that difficult. For example, recent work has suggested that increasing flux through existing pathways in C_3_ species may be enough to trigger low-level CAM under some scenarios (Bräutigam *et al.*, 2017), and comparative genomics suggest that re-wiring of pathways, rather than creating them *de novo*, likely underpins the transition to CAM (Yin *et al.*, 2018). To add to the difficulty of elucidating a mechanistic understanding of CAM evolution, the CAM phenotype is more accurately represented as a continuum, where plants can be C_3_, CAM, or a combination of both pathways called “weak” CAM (hereafter, C_3_+CAM) (Silvera *et al.*, 2010; Winter *et al.*, 2015). C_3_+CAM plants should exhibit mixed phenotypes at both physiological and genomic scales, and are potentially powerful systems for exploring the transition from C_3_ to CAM.

Our understanding of the genetics and genome structure of CAM has come predominantly from studies that involve comparisons between C_3_ and CAM tissues sampled from evolutionarily distant species, or from samples taken from one species under different age or environmental conditions (Taybi *et al.*, 2004; Cushman *et al.*, 2008*b*; Gross *et al.*, 2013; Brilhaus *et al.*, 2016) (but see (Heyduk *et al.*, 2017)). Recent studies have profiled gene expression before and after CAM induction in *Mesembryanthemum crystallinum* (Cushman *et al.*, 2008*b*) and in *Talinum* (Brilhaus *et al.*, 2016). These studies, together with comparison of transcript abundance profiles in photosynthetic (green) and non-photosynthetic (white) parts of pineapple (*Annanas comusus*) leaf blades have also provided insights into the regulation of canonical CAM genes (Zhang *et al.*, 2014; Ming *et al.*, 2015). However, RNA-seq of closely related C_3_ and CAM species, as well as intermediate C_3_+CAM lineages, are lacking.

In this study, we compared transcript profiles among three closely related *Yucca* (Agavoideae, Aspargaceae) species with contrasting photosynthetic pathways: *Y. aloifolia* consistently has nighttime uptake of CO_2_ with concomitant malic acid accumulation in leaf tissue, as well as anatomical characteristics indicative of CAM function; *Y. filamentosa* has typical C_3_ leaf anatomy and showed no positive net CO_2_ uptake or malic acid accumulation at night; *Y. gloriosa,* a hybrid species derived from *Y. aloifolia* and *Y.filamentosa*, acquires most of its CO_2_ from the atmosphere through C_3_ photosynthesis during the day with low-level CO_2_ uptake a night, but when drought stressed transitions to 100% nighttime carbon uptake (Heyduk *et al.*, 2016). *Yucca gloriosa’s* leaf anatomy is intermediate between the two parental species, and to some extent may limit the degree of CAM it can employ (Heyduk *et al.*, 2016). Clones of all three species (Supplemental Table S1) were grown in a common garden setting under both well-watered and drought stressed conditions and were sampled for gene expression and metabolomics over a 24-hour diel cycle.

## Materials and Methods

### Plant material and RNA sequencing

RNA was collected during experiments described in (Heyduk *et al.*, 2016). Briefly, clones of the three species of *Yucca* were acclimated to growth chambers with a day/night temperature of 30/17°C and 30% humidity in ∼3L pots filled with a 60:40 mix of soil:sand. Photoperiod was a 12 hour day, with lights on at 8 a.m and a light intensity of ∼380 μmol m^-2^ s^-1^ at leaf level. One clone was kept well-watered for 10 days while the second clone was subjected to drought stress via dry down beginning after the end of day 1. Clones of a genotype were randomly assigned to watered and drought treatment. On the 7^th^ day of the experiment, after plants had water withheld for the five previous days, RNA was sampled every four hours, beginning one hour after lights turned on, for a total of 6 time points; very old and very young leaves were avoided, and samples were taken from the mid-section of the leaf blade from leaves that were not shaded. Samplimg per species was biologically replicated via 3-4 genotypes per species, for a final design of 3-4 genotypes x 2 treatments x 3 species per time point sampled (Supplemental Table S1). Genotypes from the three species were randomly assigned to three different growth chamber experiments conducted in July 2014, October 2014, and February 2015. Well-watered and drought stressed clonal pairs were measured in the same experimental month. RNA was isolated from a total of 130 samples (n=36, 47, and 47 for *Y. aloifolia, Y. filamentosa,* and *Y. gloriosa*, respectively) using Qiagen’s RNeasy mini kit (www.qiagen.com). DNA was removed from RNA samples with Ambion’s Turbo DNAse, then assessed for quality on an Agilent Bioanalyzer 2100. RNA libraries were constructed using Kapa Biosystem’s stranded mRNA kit and a dual-index barcoding scheme. Libraries were quantified via qPCR then randomly combined into 4 pools of 30-36 libraries for PE75 sequencing on the NextSeq 500 at the Georgia Genomics Facility.

### Assembly and read mapping

Reads were cleaned using Trimmomatic (Bolger *et al.*, 2014) to remove adapter sequences, as well as low-quality bases and reads less than 40bp. After cleaning and retaining only paired reads, *Y. aloifolia* had 439,504,093 pairs of reads, *Y. filamentosa* had 675,702,853 pairs of reads, and *Y. gloriosa* had 668,870,164 pairs. Due to the sheer number of reads for each species, a subset was used to construct reference assemblies for each species (14% of total reads, or about 83 million read pairs per species)(Haas *et al.*, 2013). Trinity v. 2.0.6 (Grabherr *et al.*, 2011) was used for digital normalization as well as assembly. The full set of reads from each species library were mapped back to that species’ transcriptome assembly with Bowtie v. 2 (Langmead *et al.*, 2009); read mapping information was then used to calculate transcript abundance metrics in RSEM v.1.2.7 (Li and Dewey, 2011; Haas *et al.*, 2013). Trinity transcripts that had a calculated FPKM < 1 were removed, and an isoform from a component was discarded if less than 25% of the total component reads mapped to it.

To further simplify the assemblies and remove assembly artifacts and incompletely processed RNA reads, the filtered set of transcripts for each species was independently sorted into orthogroups, or inferred gene families, that were circumscribed using OrthoFinder (Emms and Kelly, 2015) clustering of 14 sequenced genomes (*Brachypodium distachyon, Phalaenopsis equestris, Oryza sativa, Musa acuminata, Asparagus officinalis, Ananas comusus, Elaeis guiensis, Acorus americanus, Sorghum bicolor, Vitis vinifera, Arabidopsis thaliana, Carica papaya, Solanum lycopersicum*, and *Amborella tricopoda)*. First, transcripts were passed through Transdecoder (http://transdecoder.github.io), which searches for open reading frames in assembled RNA-sequencing data. Transdecoder reading frame coding sequences for each species were individually matched to a protein database derived from gene models from the monocot genome dataset using BLASTx (Altschul *et al.*, 1990). Best hit for each query sequence was retained and used to sort the *Yucca* transcript into the same orthogroup as the query sequence. Assembled *Yucca* sequences were further filtered to retain only putatively full length sequences; *Yucca* transcripts that were shorter than the minimum length of an orthogroup were removed. Transdecoder produces multiple reading frames per transcript, so only the longest was retained. Scripts for orthogroup sorting and length filtering are available at www.github.com/kheyduk/RNAseq/orthoSort. Read counts for the final set of orthogrouped transcripts were re-calculated using Bowtie and RSEM and analyzed in EdgeR (Robinson *et al.*, 2010) in R 3.2.3, using TMM normalization.

To assess variation between genotypes sequenced, SNPs were calculated from the RNAseq data by mapping reads from each genotype of all species to the *Y. aloifolia* transcriptome, which is the least heterozygous of the three species. SNPs were compiled using the *mpileup* command of samtools, followed by filtering in using bcfutils. SNP positions had to have coverage between 8 and 1000, and have at least 2 alleles to be included. Indels were ignored. Similarity between genotypes and species was assessed via PCA method using the SNPRelate (Zheng *et al.*, 2012) package in R 3.2.3.

### Expression analysis of differentially expressed genes

In a given *Yucca* species, all libraries were separated by time point; within each time point, quantile adjusted conditional maximum likelihood via EdgeR (“classic mode”) was used to find the number of up and down regulated genes in response to drought stress, using a p-value cutoff of 0.05 and adjusting for multiple testing with a Holm-Bonferroni correction. Gene Ontology annotations for individual genes were obtained from the TAIR10 ontology of each gene family’s *Arabidopsis* members. GO enrichment tests were conducted for each time point per species, comparing GO categories of DE genes in well-watered vs. drought-stressed treatments, using a hypergeometric test within the phyper base function in R 3.2.3.

### Temporal profile clustering of gene expression

To assess larger patterns in the expression data while taking into account temporal patterns across time points, we employed maSigPro (Conesa *et al.*, 2006), using options for read count data (Nueda *et al.*, 2014). maSigPro is a two-step algorithm for profile clustering; the first step involves finding transcripts with non-flat time series profiles by testing generalized linear models with time and treatment factors (using a negative binomial error distribution) against a model with only an intercept (y∼1); the second step involves assessing the goodness of fit for every transcript’s regression model and assessing treatment effects. This two-step method is advantageous in that it rapidly reduces a large number of transcripts to only those that show significant variation across time, and it also readily allows users to select transcripts that have a clear expression profile (as assessed by goodness of fit of the model). For the *Yucca* data, gene regression models were initially computed on transcripts mapped per million (TMM) normalized read counts with a Bonferroni-Holm corrected significance level of 0.05. Gene models were then assessed for goodness of fit via the T.fit() function, which produces a list of influential genes whose gene models are being heavily influenced by a few data points (in this case, samples). Those genes were removed, and regression models and fit were re-calculated. Genes with significant treatment effects can either have a) different regression coefficients for the two treatments or b) different intercepts (i.e., magnitude of expression) between the two treatments. To cluster transcripts with similar profiles, we employed fuzzy clustering via the “mfuzz” option in maSigPro. The clustering steps require a user-defined value for *k* number of clusters. We assessed the optimal number of clusters for each species’ data by examining the within group sum of squares for *k*=1:20 clusters. A *k* was chosen where the plot has a bend or elbow, typically just before the group sum of squares levels off for higher values of *k.* A *k* of 9, 12, and 15 was used for *Y. aloifolia, Y. gloriosa*, and *Y. filamentosa*, respectively. To estimate *m*, the “fuzzification parameter” for fuzzy clustering, we employed the mestimate() function in the Mfuzz package. *m* of 1.06, 1.05, and 1.05 was estimated for *Y. aloifolia*, *Y. gloriosa*, and *Y. filamentosa.*

By default, the clustering steps in maSigPro are run on genes that are not only significant with regards to temporal expression (non-flat profiles across time), but also only on the subset of genes that are significantly different between treatments. We modified the code for the see.genes() function to fuzzy cluster all transcripts that had non-flat profiles, regardless of whether they showed a significant change in expression as a result of drought stress. Afterwards, we found transcripts that were significantly different between treatments with an R-squared cut off of 0.7. The modified code for the see.genes() function, as well as detailed guide to the steps taken for this analysis, are available at www.github.com/kheyduk/RNAseq.

For genes of interest, additional tests were done to assess whether species differed significantly in their temporal pattern of expression. Because count data cannot be accurately compared between species, we instead used TPM values. For each gene family of interest, we selected a single transcript per species, typically one that was highest expressed and had time-structured expression as determined by maSigPro. TPM values were scaled within each species’ transcript, separately for well-watered and drought-stressed libraries, by the maximum TPM value. All TPM values for each gene for each treatment had a polynomial model fit with degree=5 without distinguishing species, and a second polynomial model that included species as a factor. Using ANOVA, we compared the fits of the two models. For genes that had a significantly better fit when species was treated as a factor, we report the t-statistic and p-value for the species that had a significant coefficient.

### Gene annotation and gene tree estimation

All transcripts were first annotated by their membership in gene families; gene family annotations were based on the *Arabidopsis* sequences that belong to the gene family, using TAIR10 annotations. To address homology of transcripts across species, gene trees were constructed from protein-coding sequences of gene families of interest, and included *Yucca* transcript sequences as well as the 14 angiosperm sequenced genomes. Nucleotide sequences were first aligned via PASTA (Mirarab *et al.*, 2014). Gene trees were estimated via RAxML (Stamatakis, 2006) using 200 bootstrap replicates and GTRGAMMA nucleotide model of substitution.

### Metabolomics

Samples for starch and metabolomics were collected from a separate experiment conducted in February 2017 at the University of Georgia greenhouses. Growth conditions in the chamber were identical to conditions used when harvesting tissue for RNA-Seq (above), and plants used were the same genotype, but not the same clone, as for RNA-Seq. As this was only a preliminary analysis, samples for starch and metabolites were collected only for the parental species, and only under well-watered conditions. Samples were collected every 4 hours starting 1 hour after the lights turned on from 6 replicate plants per species; replicates were from different genotypic backgrounds (see Supplemental Table S1). Gas exchange data was collected concurrently to ensure plants were behaving as when RNA was collected previously (Supplemental Figure S1). Tissue for starch was flash frozen in liquid N2, then later dried in a forced air oven. Samples were ground and 0.02 g was washed first with room temperature acetone, then with 80% EtOH, and finally heated at 90C in 1% hydrochloric acid and centrifuged to pellet any remaining tissue (Hansen and Moller, 1975; Oren *et al.*, 1988). A 1:40 dilution of 5% Lugol’s idodine was added to starch extracts and measured in a spectrophotometer at 580 nm. Values were compared against a standard curve made from corn starch dissolved in 1% hydrochloric acid.

Tissue for metabolic analysis was flash frozen in liquid N_2_ then stored at −80°C until samples were freeze-dried. A 1:1 mixture of MeOH and chloroform (400μL) was added to 10mg of freeze-dried, ball-milled (Mini-beadbeater, Biospec products Bartlesville OK, USA) tissue along with adonitol as an internal standard. Mixtures were sonicated for 30 minutes at 8-10°C, equilibrated to room temperature, and polar metabolites recovered by liquid phase partitioning after 200μL H_2_O was added to the extract. Ten μL of the aqueous-methanol phase was dried and derivatized for GCMS by adding 15μL methoxyamine hydrochloride and incubating at 30°C for 30 minutes, then by adding 30μL MSTFA and incubating at 60°C for 90 minutes. Derivatized samples were analyzed via gas chromatography as in Frost et al (2012). Chromatograms were deconvoluted using AnalyzerPro (SpectralWorks, Runcom, UK). Peak identities were based on NIST08, Fiehnlib (Agilent Technologies, (Kind *et al.*, 2009)), and in-house mass spectral libraries. Peak matching between samples was based on the best library match according to AnalyzerPro and retention index (Jeong *et al.*, 2004). Initial metabolite peak calls were filtered first by the confidence level of their best library match (>0.5) and then by raw peak area (>1000). Filtered metabolite peak areas were then normalized based on adonitol peak areas. Standard curves were run for ascorbate, sucrose, malic acid, and citric acid to determine absolute concentrations in umol/g of dry weight (Supplemental Figure S5).

Normalized values were imported into R v. 3.3.3 and, where appropriate, multiple metabolite peaks were summed to obtain a single value per metabolite. Time points 3 and 6 (last day time point and last night time point) were removed from analysis due to errors in derivitization steps. Remaining values were filtered for sample presence, retaining only metabolites that were found in at least 25% of all samples. The resulting 217 metabolites were imported into maSigPro, where we tested for time-structured expression using species as a treatment in the design matrix, allowing for polynomials with degree=3, and using a quasipoisson distribution in the glm model.

Data generated is available on NCBI’s Short Read Archive (RNA-seq data, BioProject #PRJNA413947), or at github.com/karohey/RNAseq_Yucca/C_3_-CAM (for count matrices and metabolite data).

## Results

### Photosynthetic phenotypes

As described in previous work, *Y. aloifolia* conducts atmospheric CO_2_ fixation at night via CAM photosynthesis, while *Y. filamentosa* relies only on daytime CO_2_ fixation and the C_3_ cycle. *Yucca gloriosa*, a C_3_-CAM intermediate species, uses mostly daytime CO_2_ fixation with low levels of nocturnal gas exchange under well-watered conditions, then relies solely on CAM photosynthesis under drought stress (Fig. 1B). Gas exchange and titratable acidity measurements shown in Fig. 1 are from prior work when RNA was sampled, though gas exchange patterns were largely consistent in *Y. aloifolia* and *Y. filamentosa* during a second round of sampling for metabolites (Supplemental Figure S1).

### Assembly and differential expression

After filtering to remove low abundance transcripts (FPKM<1) and minor isoforms (<25% total component expression), an average of 55k assembled transcripts remained per species. Transcripts were then sorted into gene families (orthogroups) circumscribed by 14 sequenced plant genomes and removed if their length was shorter than the minimum length for a gene family. Considering only transcripts that sorted into a gene family and had the proper length, transcriptome sizes were reduced further: 19,399, 23,645, and 22,086 assembled transcripts remained in *Y. aloifolia, Y. filamentosa,* and *Y. gloriosa,* respectively. SNPs showed greater variation between species rather than among genotypes (Fig. 2A), although *Y. filamentosa* exhibited more SNP variation among genotypes than the other two species. *Yucca gloriosa* genotypes used in this study were found to be slightly more similar to *Y. aloifolia* than *Y. filamentosa* based on PCA analysis of SNP distances; this is likely a consequence of choosing *Y. gloriosa* accessions that showed a propensity for CAM (and therefore were potentially more similar to *Y. aloifolia*) under drought stress for RNA-seq.

**Figure 2.**
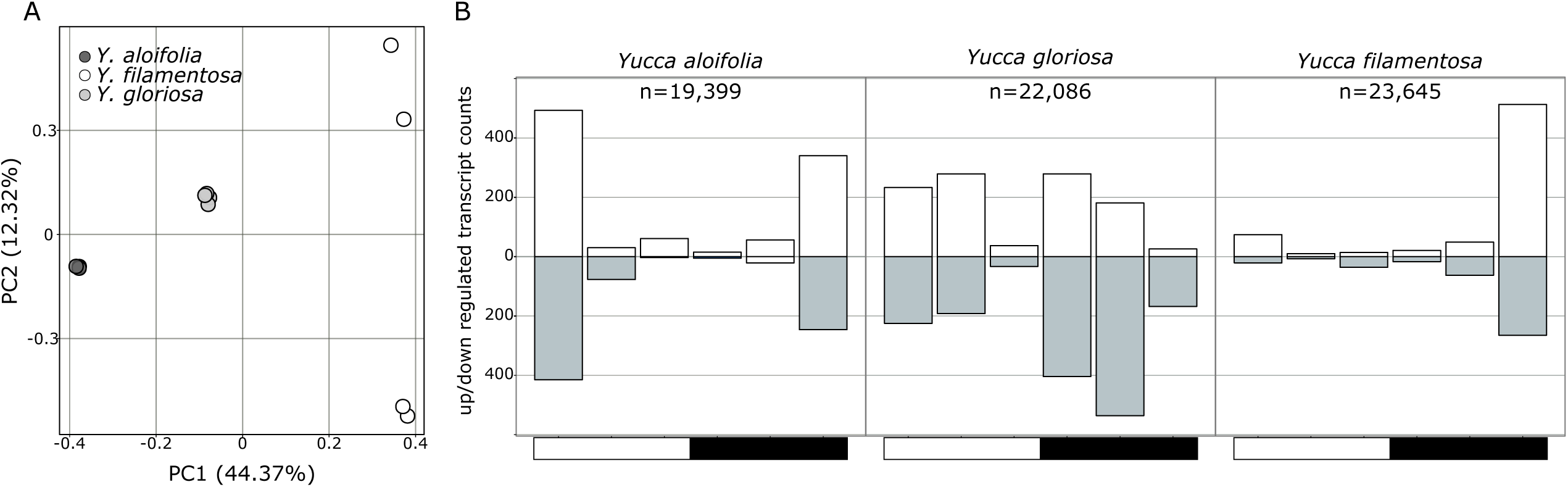
Overview of genetic diversity and differential gene expression across Yucca. A) PCA of SNP diversity from transcriptome data, and B) up/down differential expression between well-watered and drought stressed plants at each time point based on EdgeR, with counts as a proportion of total transcripts expressed.

Differential expression analysis at each time point between well-watered and drought-stressed samples showed distinct patterns in the three species (Fig. 2B). The effect of drought on expression was greatest one hour after the start of the light period in the CAM species *Yucca aloifolia,* but just before light in the C_3_ species *Y. filamentosa*. *Yucca gloriosa* (C_3_-CAM intermediate) had near constant levels of differentially expressed transcripts across the entire day/night cycle. Gene Ontology (GO) enrichment tests showed general processes, such as metabolism and photosynthesis, as being commonly enriched in the differentially expressed transcripts (Supplemental Table S2).

Transcripts of each species were classified as time structured if their expression across time under either well-watered and drought conditions could be better described with a polynomial regression, rather than a flat line, with significance-of-fit corrected for multiple tests. There were 612, 749, and 635 transcription factor annotated transcripts with time-structured expression profiles in *Y. aloifolia*, *Y. filamentosa,* and *Y. gloriosa*, respectively. Of those, 92, 62, and 83 were differentially expressed in *Y. aloifolia, Y. filamentosa,* and *Y. gloriosa,* respectively, under drought (Supplemental Table S3). Putative CAM pathway genes (Fig. 1) largely showed the expected expression patterns in *Y. aloifolia*, and additionally all three species shared time-structured gene expression patterns for some canonical CAM genes regardless of photosynthetic pathway. In all three *Yuccas*, PEPC, its kinase PPCK (Fig. 3A and B), as well as decarboxylating enzymes NAD/P-me, PPDK and, PEPCK showed time-structured expression (Supplemental Figure S2). PEPC and PPCK (Fig. 3A and B) exhibited time-structured expression in all three species, though they were only differentially expressed between well-watered and droughted treatments in *Y. gloriosa*. Expression of PEPC in *Y. filamentosa* was much lower in terms of TPM (transcripts per kilobase million), but had the same temporal pattern as both *Y. aloifolia* and *Y. gloriosa* (well-watered: F_(12,46)_ = 1.38, p < 0.212; drought: F_(12,42)_=0.53, p < 0.866). In all 3 species, PPCK was most highly expressed at night (Fig. 3B), consistent with its role in activating PEPC for dark carboxylation, and showed no difference in temporal expression across species (well-watered: F_(12,46)_ = 0.85, p < 0.605; drought: F_(12,42)_=1.21, p < 0.309). Carbonic anhydrase (CA), involved in conversion of CO_2_ to HCO_3_-, had only 3 transcripts that were temporally structured in their expression in *Y. aloifolia*; two α-CA and one γ. In none of these cases did expression increase at night as might be expected (Supplemental Figure S3).

**Figure 3.**
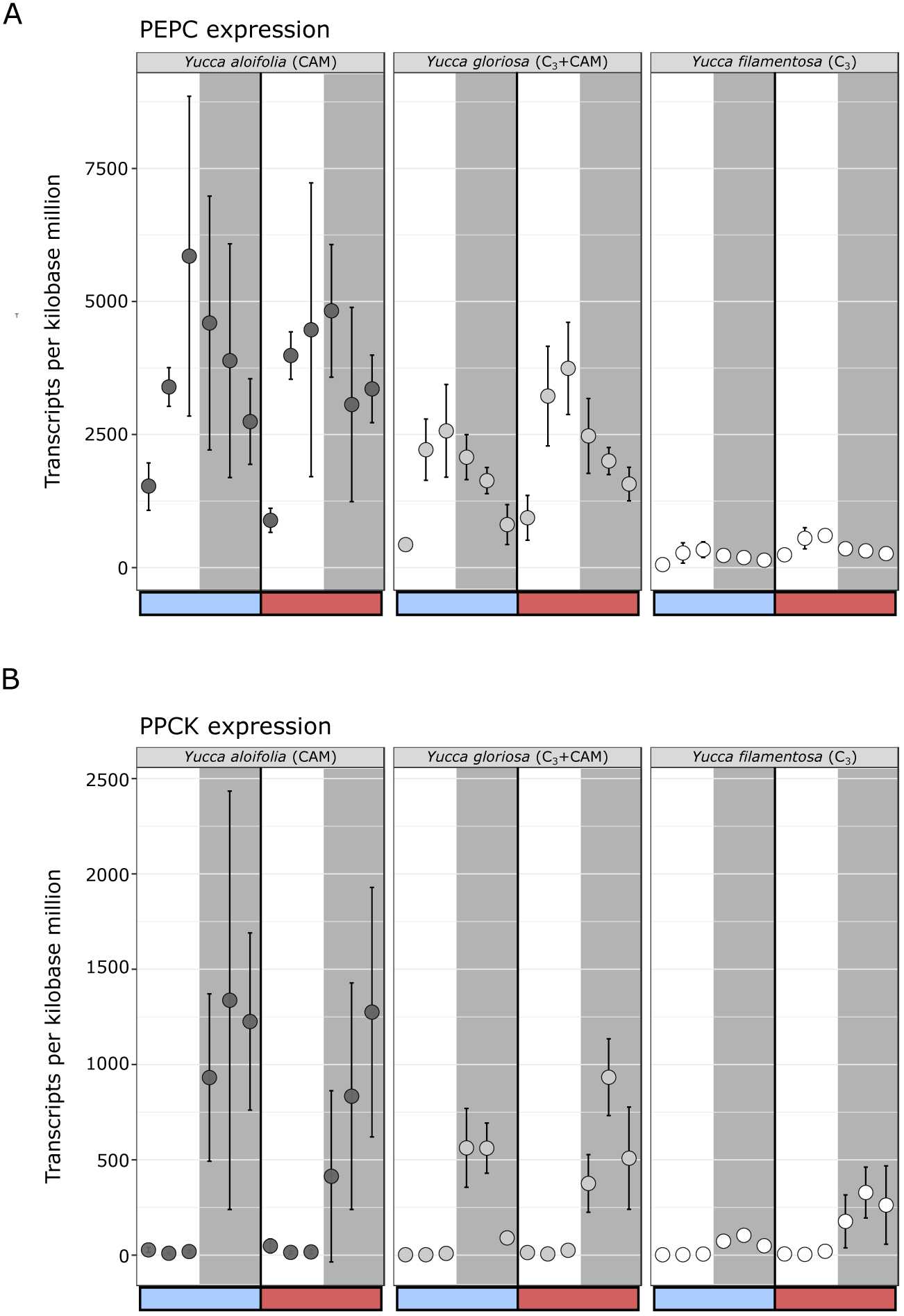
Gene expression for PEPC (A) and PPCK (B) in all three *Yucca* species, shown for day (white background) and night (grey background) time points, under both well-watered (blue bar) and drought-stressed (red bar) conditions. Mean TPM ± one standard deviation is plotted. Transcripts shown represent all copies in each gene family (note, an alternative, but more lowly expressed gene family exists for PEPC and is not shown here), and all transcripts were significantly time-structured in all three species.

### Metabolomics

Of the 214 metabolites that were present in at least 25% of samples, 87 had a significant fit to a polynomial regression line (Fig. 4), with 16 having significant differences in either abundance or temporal regulation between the *Y. aloifolia* and *Y. filamentosa* (R^2^>0.5) (Supplemental Table S4). Starch degradation is one possible route CAM plants can use for the nightly regeneration of PEP. Whilst starch content overall was comparable between the C_3_ and CAM species, there was no net dark depletion of starch in the CAM species, suggesting little reliance on starch for nocturnal generation of PEP in the CAM Yucca (Fig. 5A). In contrast, starch is degraded at night in the C_3_ species and hybrid, with increased levels of α-glucan phosphorylase (PHS), a gene responsible for phosphorolytic degradation of starch (Smith *et al.*, 2005; Borland *et al.*, 2016). Maltose, a starch-derived breakdown product, was substantially elevated in the C_3_ species compared to the CAM (Fig. 5B). The difference in maltose content was reflected by higher expression of the maltose exporter MEX1 gene in *Y. filamentosa* (Fig. 5B). Malic acid had greater turnover in *Y. aloifolia,* and transcript abundance of malate dehydrogenase (MDH), responsible for interconversion of malic acid and oxaloacetate (Fig. 5C), was likewise higher in the CAM species.

**Figure 4.**
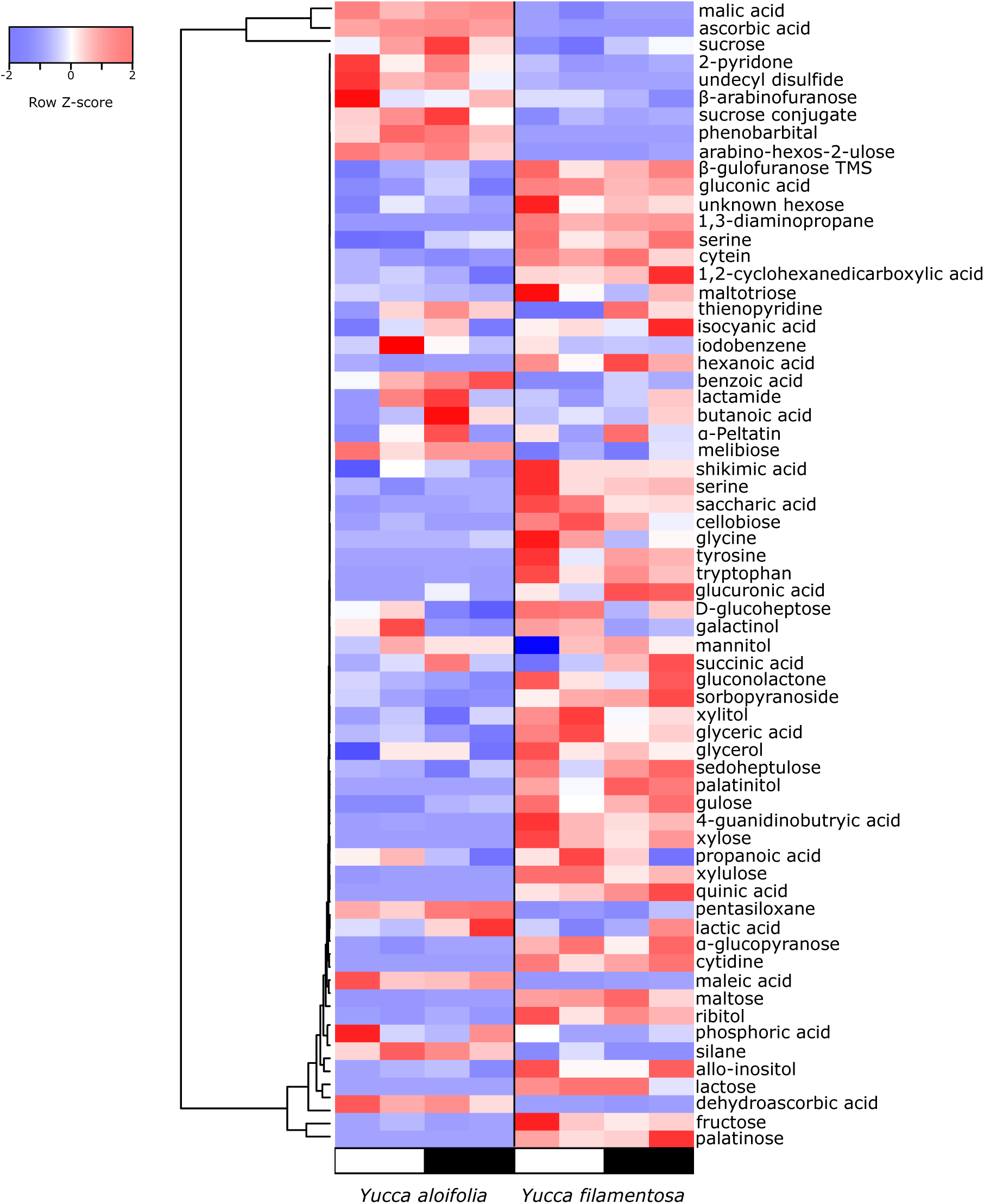
Heatmap of abundance of a biologically meaningful subset of metabolites that had time-structured fluctuations in abundance (i.e., could be fit to a polynomial), shown for each species during the day (white bar) and night (black bar).

**Figure 5.**
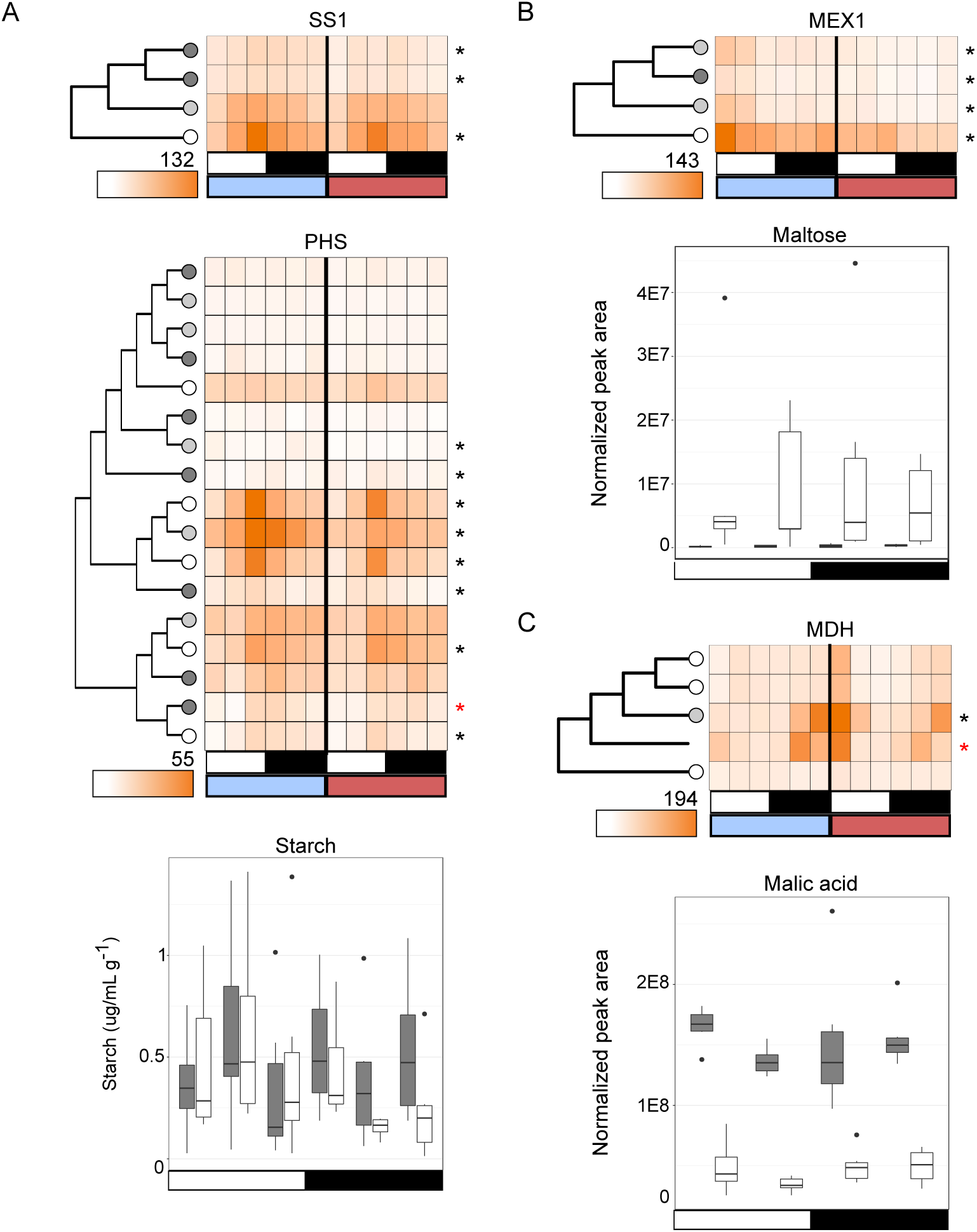
Gene expression and related metabolites, shown over a day (white bar) and night (black bar) period, under both well-watered (blue bar) and drought-stress (red bar) in RNA-seq data only. Maximum TPM for each gene shown below the gene tree; asterisks indicate transcripts that were significantly time-structured, with red coloring indicating differential expression between watered and drought. Gene tree circles are color coded by species (dark grey=Y. aloifolia (CAM), white=Y. filamentosa (C3), light grey=Y. gloriosa (C3-CAM). The colors are carried to the metabolite plots (dark grey bars=Y. aloifolia, white bars=Y. filamentosa). A) Starch synthase 1 (SS1), involved in the production of starch; glucan phosphorylase (PHS), involved in degradation of starch B) Maltose exporter 1 (MEX1), transports maltose out of plastids. C) Malate dehydrogenase (MDH), responsible for interconversion of oxaloacetate and malic acid.

An alternative source of carbohydrates for PEP can come from soluble sugars. Several soluble sugars had higher abundance in *Y. filamentosa*, including fructose and glucose (Fig. 6A). Fructose (but not glucose or sucrose) had a significant temporal difference between *Y. aloifolia* and *Y. filamentosa* (Supplemental Figure S4), with concentrations in *Y. filamentosa* decreasing during the dark period while concentrations in *Y. aloifolia* remained flat. Both species accumulate similar amounts of sucrose (Fig. 6A), indicating no difference in the amount of hexoses dedicated to sucrose production. There is a slight temporal change across the day-night period in *Y. aloifolia,* but it was not significant based on polynomial regression analysis. Gene expression also does not implicate conversion of hexose to triose phosphates as a mechanism for generating differences in hexose concentrations: both *Y. aloifolia* and *Y. filamentosa* express fructose 1,6-bisphosphate aldolase (FBA) at equal levels, although different gene copies are used in *Y. aloifolia* vs. *Y. filamentosa* (Fig. 6A), and *Y. filamentosa* has a significantly different timing of expression relative to *Y. aloifolia* under both well-watered (F_(12,46)_ = −4.293, p = 8.99e-05) and drought-stressed conditions (F_(12,42)_ = −6.79, p = 3.55e-08). The different FBA paralogs expressed in *Y. aloifoia* and *Y. gloriosa* compared to *Y. filamentosa* represent alternative localizations; the FBA homolog expressed in *Y. filamentosa* has an *Arabidopsis* ortholog which localizes to the chloroplast, while the copy expressed in *Y. aloifolia* and *Y. gloriosa* has cytosolic *Arabidopsis* ortholog. FBA in the chloroplast is responsible for the production of metabolites for starch synthesis, implicating starch synthesis and breakdown in *Y. filamentosa*, consistent with this species’ increase in maltose production. The cytosolic version found in the CAM and C_3_+CAM intermediate species is thought to be involved in glycolysis and gluconeogenesis.

**Figure 6.**
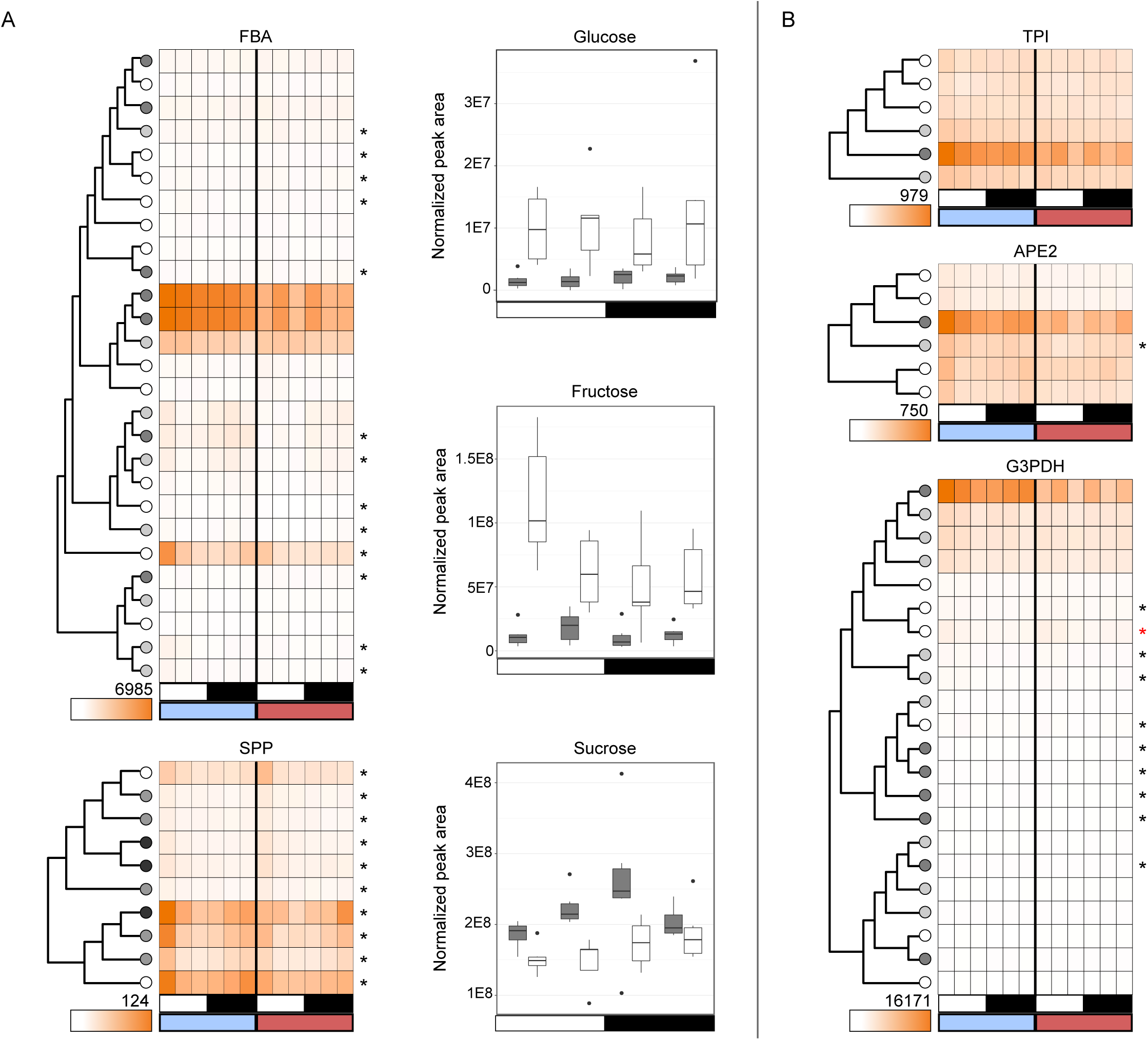
Gene expression and related metabolites, shown over a day (white bar) and night (black bar) period, under both well-watered (blue bar) and drought-stress (red bar) in RNA-seq data only. Maximum TPM for each gene shown below the gene tree; asterisks indicate transcripts that were significantly time-structured, with red coloring indicating differential expression between watered and drought. Gene tree circles are color coded by species (dark grey=Y. aloifolia (CAM), white=Y. filamentosa (C3), light grey=Y. gloriosa (C3-CAM). The colors are carried to the metabolite plots (dark grey bars=Y. aloifolia, white bars=Y. filamentosa). A) Fructose bisphosphate aldolase, responsible for interconversion of fructose-6-P and triose phosphates, and sucrose phosphatase (SPP) produces sucrose from glucose and fructose molecules. B) Triose phosphate isomerase (TPI) interconverts the two forms of triose phosphates, APE2 is a triose phosphate transporter out of the plastid, and G3PDH is involved glycolysis.

Triose phosphates are too small to measure through GC-MS metabolomics methods, but genes associated with interconversion of glyceraldehyde 3-phosphate and dihydroxyacetone phosphate (triose phosphate isomerase, TPI) as well as genes involved in transport (triose phosphate transporter, APE2) show higher expression in *Y. aloifolia* compared to both *Y. filamentosa* and *Y. gloriosa* (Fig. 6B), though genes do not significantly differ in temporal expression pattern based on post-hoc linear model tests. G3PDH (glyceraldehyde-3-phosphate dehydrogenase), an enzyme involved in downstream branches of glycolysis, likewise has highest expression in *Y. aloifolia;* accounting for species in the linear model of gene expression significantly increases fit of the model under both well-watered (F_(12,46)_ = 2.45, p < 0.015) and drought-stressed conditions (F_(12, 42)_ = 4.94, p < 4.97e-5).

Metabolites involved in reactive oxygen species (ROS) scavenging pathways also showed large differences between *Y. aloifolia* and *Y. filamentosa*. Vitamin C, or ascorbic acid, was present at much higher levels in *Y. aloifolia* (Fig. 7A), as was the oxidized form dehydroascorbic acid (Fig. 7B). Neither had a temporal expression pattern, however, indicating constant levels of both metabolites across the day-night period. Previous work has implicated increases in ascorbic acid as a method for CAM plants to remove ROS produced by high nocturnal respiration (Abraham *et al.*, 2016), but genes involved in mitochondrial respiration (cytochrome-C, CYTC) are only slightly elevated in *Y. aloifolia* at night (Fig. 7C). The biogenesis of ascorbate through galactono-gamma-lactone dehydrogenase (GLDH) does not seem to differ between the two parental *Yucca* species either, based on gene expression (Fig. 7C).

**Figure 7.**
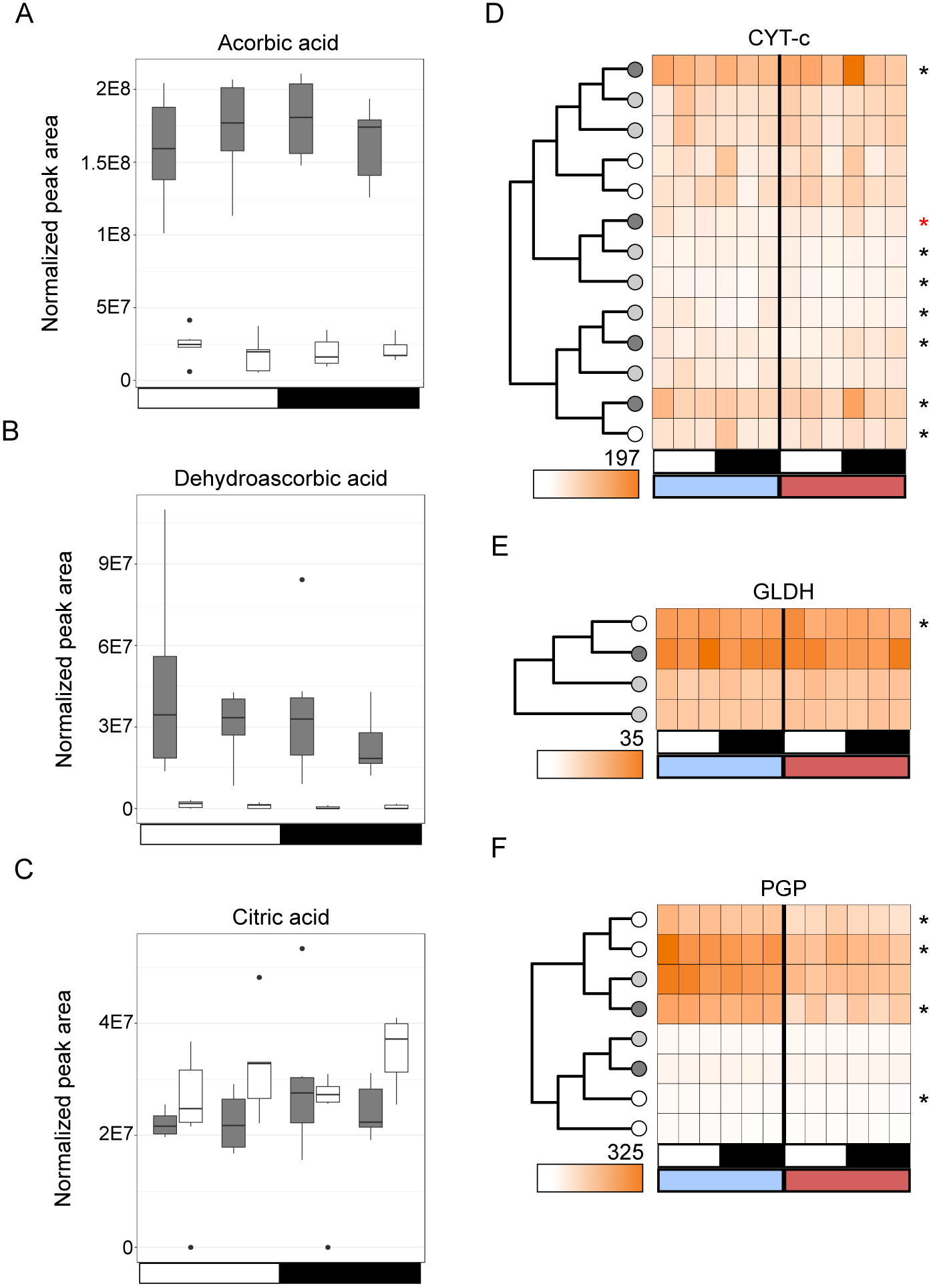
Abundance over the day (white bar) and night (black bar) period for A) ascorbic acid, B) dehydroascorbic acid, and C) citric acid. Gene expression for D) cytochrome-C (CYTC), E) galactono-gamma-lactone dehydrogenase (GLDH), and F) phosphoglycolate phosphatase (PGP) over the day (white bar) and night (black bar), under both well-watered (blue bar) and drought-stressed (red bar) conditions. Maximum TPM for each gene shown below the gene tree; asterisks indicate transcripts that were significantly time-structured, with red coloring indicating differential expression between watered and drought. Gene tree circles are color coded by species (dark grey=Y. aloifolia (CAM), white=Y. filamentosa (C3), light grey=Y. gloriosa (C3-CAM). The colors are carried to the metabolite plots (dark grey bars=Y. aloifolia, white bars=Y. filamentosa).

Alternatively ROS might be produced from daytime activities, however whether or not CAM plants reduce oxidative stress (via reduced photoihibition, (Adams and Osmond, 1988; Griffiths *et al.*, 1989; Pieters *et al.*, 2003)) or instead produce high levels of O_2_ (from increased electron transport behind closed stomata, (Niewiadomska and Borland, 2008)) remains unclear in the literature. Regardless, the photorespiratory pathway has been proposed as a means of protection from oxidative stress (Kozaki and Takeba, 1996), but reduced photorespiration is thought to be a key benefit of CAM photosynthesis. Kinetic modeling of RuBisCO oxygenase/carboxylase activity suggested CAM plants have either equivalent levels of photorespiration to C_3_ species or reduced by as much as 60% (Lüttge, 2010). An increase in antioxidants like ascorbic acid would therefore be beneficial in CAM plants if photorespiration is reduced relative to C_3_ while O_2_ and ROS are still produced at the same rate. In support of this hypothesis, phosphoglycolate phosphatase (PGP) – a gene involved in the first step in breaking down the photorespiratory product 2-phosphoglycolate – is elevated in *Y. filementosa* and *Y. gloriosa* relative to the CAM species (Fig. 7F). Metabolites that take part in photorespiration, including glycine and serine, peak in the day period in C_3_ *Y. filamentosa*, and are generally much higher in *Y. filamentosa* than seen in *Y. aloifolia* (Fig. 4) (Scheible *et al.*, 2000; Novitskaya *et al.*, 2002), suggesting CAM *Y. aloifolia* may instead rely on increased ascorbic acid concentrations to reduce ROS stress.

## Discussion

### CAM pathway genes

Time-structured expression of key CAM genes in a C_3_ species of *Yucca* suggests ancestral expression patterns required for CAM may have predated its origin in *Yucca*. This important observation is in line with recent suggestions that the frequent emergences of CAM from C_3_ photosynthesis was facilitated by evolution acting directly on a low flux pathway already in place for amino acid metabolism (Bräutigam *et al.*, 2017). Flux analysis using ^13^C labelled substrates has shown that C_3_ plants can use organic acids at night to fuel amino acid synthesis (Gauthier *et al.*, 2010; Szecowka *et al.*, 2013). Such carbon labeling experiments, in addition to shared gene expression shown here, suggest that the evolution of CAM may simply require increasing the flux capacity of existing carboxylation pathways in C_3_ plants, without the need for extensive re-wiring or diel rescheduling of enzymes (Bräutigam *et al.*, 2017).

### Carbohydrate metabolism

To provide the nightly supply of PEP needed as substrate for CO_2_ and PEPC, CAM plants either break down soluble sugars (including polymers of fructose in fructans) or starches to regenerate PEP via glycolysis. Work in the closely related genus *Agave* indicates that soluble sugars are the main pool for nightly PEP regeneration (Abraham *et al.*, 2016). As seen in *Agave*, the CAM *Yucca* species use soluble sugars as a carbohydrate reserve for PEP requirements, while C_3_ *Y. filamentosa* likely relies on starch pools. Although starch concentrations were largely equal in *Y. aloifolia* and *Y. filamentosa*, degradation of starch to form maltose was significantly higher in *Y. filamentosa*. The low levels of MEX1 and PHS1 expression in *Y. aloifolia* further suggests that starch degradation was recently lost (or gained in *Y. filamentosa*) as *Y. aloifolia* and *Y. filamentosa* diverged only 5-8MYA (Good-Avila *et al.*, 2006; Smith *et al.*, 2008). *Yucca gloriosa* has intermediate expression of MEX1 and PHS relative to its parental species, indicating some reliance on starch for carbohydrates like its C_3_ parent.

Soluble sugars, such as glucose, fructose, and sucrose, can serve as an alternative source of carbohydrates for glycolysis. In *Agave*, fructans (chains of fructose monomers) are the predominant source of nocturnal carbohydrates for PEP (Wang and Nobel, 1998; Arrizon *et al.*, 2010). *Agave*, relative to *Arabidopsis*, has temporal regulation of soluble sugar production and a 10-fold increase in abundance (Abraham *et al.*, 2016). In general, there was a lack of diel turnover in soluble sugars in *Y. aloifolia*, although it is possible unmeasured fructans constitute the majority of the carbohydrate pool. With one exception neither species shows temporal fluctuation of abundance of soluble sugars (*Y. filamentosa* exhibits time-structured variation in fructose concentrations, Supplemental Figure S4). Sucrose concentrations are largely equal between the two species, while glucose and fructose are elevated in C_3_ *Y. filamentosa*. Glucose and fructose are the building blocks of sucrose, but it is unclear from the metabolite and transcript data alone whether these are elevated in *Y. filamentosa* due to degradation of sucrose, or for some other purpose.

Many of the genes involved in glycolytic processes had much higher expression in *Y. aloifolia*, suggesting that the breakdown of triose phosphates into PEP is occurring at a higher rate in CAM *Yuccas*. Fructose bisphosphate aldolase (FBA) acts as a major control point for glycolysis by converting fructose 1,6-bisphosphate into triose phosphates and is also involved in the reverse reaction in the Calvin Cycle (formation of hexose from triose phosphates). FBA expression is initially high in both parental species (Fig. 6A), then rapidly drops in the C_3_ species and is sustained throughout the day period in both *Y. aloifolia* and *Y. gloriosa,* although alternative copies of this gene are used in CAM and C_3_ parental species. FBA is thought to be driven toward triose phosphate production within the cytosol (the gene copy expressed in the CAM species), whereas the chloroplastic copy expressed in the C_3_ species is involved in Calvin Cycle carbohydrate synthesis. Gene expression patterns therefore suggest that while *Y. aloifolia* expresses FBA for production of triose phosphates for glycolysis and PEP regeneration, *Y. filamentosa* uses the reverse reaction to synthesize greater concentrations of soluble sugars.

In total, metabolite data and gene expression suggest soluble sugar pools in and of themselves are not the critical part of carbon metabolism for CAM in *Yucca*; instead, it is more likely that flux through the system, particularly through glycolysis, is important for the maintenance of PEP and thus effective CAM function. The apparent variation in *which* carbohydrate pool is used – starch for C_3_, soluble sugars for CAM – is surprising, given the relatively short evolutionary distance between the two species. The functional importance of large glucose and fructose accumulation and retention in *Y. filamentosa* relative to *Y. aloifolia* is unclear. Roles for the hexoses glucose and fructose in C_3_ plants include hormonal signaling (Zhou *et al.*, 1998; Arenas-Huertero *et al.*, 2000; Leon and Sheen, 2003), plant growth and development (Miller and Chourey, 1992; Weber *et al.*, 1997), and gene expression regulation (Koch, 1996); because CAM plants undergo all of the same metabolic processes, the stark difference in concentrations of these hexoses in C_3_ and CAM *Yucca* remains to be investigated. Metabolite data presented in this study is preliminary, and was meant to highlight differences in metabolite pools between these closely related C3 and CAM species. Further studies to describe the parental C_3_ and CAM species metabolomes behave under drought stress, as well as the metabolic profile of the C_3_-CAM *Yucca* hybrid, will provide a greater understanding for the links between metabolites, carbon metabolism, and photosynthesis.

### Antioxidant response in CAM

Previous work in *Agave* discovered high levels of ascorbate and NADH activity relative to C_3_ *Arabidopsis*, which was thought to be due to increases in mitochondrial activity at night in CAM species relative to C_3_ (Abraham *et al.*, 2016). Similarly, *Yucca aloifolia* has much higher levels of ascorbic acid and dehydroascorbic acid relative to its C_3_ sister species and implies different requirements for antioxidant response between the two species. For example, respiration rates might be expected to be higher in CAM species at night to sustain the active metabolism. Although citric acid abundance is nearly identical in C_3_ and CAM *Yucca* species, expression of cytochrome-C, a part of the mitochondrial electron transport chain, is higher in the CAM *Y. aloifolia*. Alternatively, due to inhibited photorespiratory response in the CAM species, an alternative form of ROS scavenging may be needed to regulate oxidation in the cells resulting from either photoinhibition or O_2_ accumulation from electron transport behind closed stomata during the day. It is possible CAM species are using antioxidant metabolites like ascorbic acid to prevent oxidative stress, rather than relying on photorespiration. Indeed genes (PGP) and metabolites (glycine and serine) involved in photorespiration were more lowly expressed and found in lower abundance, respectively, relative to C_3_ *Y. filamentosa.* Whether or not increased antioxidant response is required for CAM to efficiently function in plants is unknown, and future work discerning ROS production and mitigation – particularly in the hybrid *Y. gloriosa* – will inform understanding of the role of ROS scavenging and its impact on photosynthetic functions.

Transcriptomics and metabolomics of the parental species *Y. aloifolia* and *Y. filamentosa* revealed many changes to regulation, expression, and abundance. The most notable differences included degree of expression of core CAM genes and fundamental differences between the C_3_ and CAM species in starch and soluble sugar metabolism. The increased reliance on soluble sugars in the CAM species, which is not shared with the C_3_ *Y. filamentosa,* indicates a recent alteration to carbohydrate metabolism after the divergence of these two species and coincident with the origin of the CAM pathway. The diploid hybrid species, *Y. gloriosa*, exhibited gene expression profiles more similar to its CAM parent, *Y. aloifolia*, than the C_3_ parent, *Y. filamentosa.* Additionally, the CAM species *Y. aloifolia* had heightened antioxidant response (both in metabolites and gene expression) relative to *Y. filamentosa*, indicating that the operation of CAM imposes a significant oxidative burden. Despite these differences, similarities exist in levels of gene expression of a few CAM genes (PEPC, for example) between the C_3_ and CAM *Yuccas* studied here, perhaps indicated shared traits in an ancestral genome that may have facilitated the convergent evolution of CAM photosynthesis within the Agavoideae. Continued comparative research on closely related C3 and CAM species, as well as intermediate forms, is necessary to understand the genetic underpinnings of CAM, as well as to determine whether ancestral genetic enabling has facilitated the evolution of CAM.

## Supplemental data

Table S1 - Genotypes sequenced through RNAseq

Figure S1 - Gas exchange data for metabolomics samples.

Table S2 - GO term enrichment

Table S3 - Differentially 496 expressed transcription factors

Figure S2 - Decarboxylation gene expression

Figure S3 - Carbonic anhydrases gene expression

Table S4 - Significantly different metabolites between C3 and CAM Yucca.

Figure S4 - Temporally variable metabolite regressions

Figure S5 - Calibrated concentrations of key metabolites

## Acknowledgements

Authors would like to acknowledge support from the University of Georgia and the National Science Foundation (DEB 1442199).

